# Enhancing fMRI language lateralization indices by homotopic mapping

**DOI:** 10.1101/792515

**Authors:** Matteo Pesci, Mariska van Steensel, Nick Ramsey, Mathijs Raemaekers

**Affiliations:** University Medical Center Utrecht

## Abstract

The non-invasive nature of fMRI would make it an excellent tool for determining language lateralization, potentially replacing the WADA test that is now considered the gold standard. However, while the result of the WADA test has 3 straightforward categories, i.e. left, right, or bilateral language lateralization, fMRI lateralization indices (LIs) are on a continuous scale between −1 (completely right lateralized) and 1 (completely left lateralized), requiring arbitrary thresholding to come to a useful clinical measure. We hypothesize that the absence of clear categories in fMRI LIs is in part linked to imperfect fMRI task control and/or homotopically mapped language activity in the non-dominant hemisphere. In this study we attempt to account for these sources of brain activity by making a detailed homotopic comparison between voxels in the language areas and their homologues, which is now possible using the Cgrid toolbox. 36 epilepsy were included who underwent presurgical fMRI including a picture naming and verb generation task, in addition to the WADA test, functional Transcranial Doppler sonography (fTCD), or Electrocortical stimulation (ECS). LIs were calculated using a traditional approach, and a novel approach based on the paired comparison between language voxels and their homologues. We observed that lateralization indices came in closer alignment with those established by the WADA test, ECS, of fTCD. Activity maps indicated that the accounting for imperfect fMRI task control was at least part of the reason for the improvement when using the new approach. This Cgrid based approach could help fMRI replacing the more invasive WADA test in the future.

## Introduction

To reduce the risk of postoperative cognitive deficits, lateralization of language function has to be assessed accurately prior to surgery. In healthy individuals, language function is lateralized to the left hemisphere in 73%–96% of cases (Springer et al. 1999; Knecht et al. 2000), but atypical representation of language (bilateral or right-dominant) might occur more frequently in congenital neurological conditions, e.g. certain cases of epilepsy (Springer et al. 1999).

For testing language lateralization, the Wada test (intracarotid amobarbital test) is considered the gold standard for preoperative assessment of lateralization of language (Wada and Rasmussen 2007). During the WADA test amobarbital is injected in the internal carotid artery, which causes functional disruption of the ipsilateral hemisphere, and temporary loss of language function dependent on hemispheric dominance. Alternatively, language lateralization can also be tested using electrocortical stimulation (ECS) (Wyllie et al. 1990), where a current is produced between two adjacent electrodes of a subdural grid, temporally disrupting the functioning of the cortex beneath. Both of these approaches are however highly invasive and can cause major distress to the patients.

Alternatively, functional magnetic resonance imaging (fMRI) is one of the emergent techniques that might offer a safe, non-invasive and relatively rapid alternative to the Wada test or ECS (Fernández et al. 2001; Medina et al. 2004, 2005). The output of fMRI is however often less straightforward to interpret. While the outcome of e.g. the Wada test is trichotomous: left, right or bilateral representation of language, the outcome of fMRI determined language lateralization is typically a lateralization index (LI) which has a continuous scale. The fMRI LI is based on the number of left vs. right activated voxels during a language task and ranges from 1 (fully left lateralized) to −1 (fully right lateralized). As the fMRI-derived has LI to be converted into a trichotomous and clinically useful measure, arbitrary decisions have to be made. The objective of this study is to bring language lateralization according to fMRI and those established by the Wada test or ECS in closer alignment, making such decisions less arbitrary.

One possible explanation for incomplete language lateralization in fMRI is that it is effectively impossible to design a language-paradigm that fully isolates language specialized voxels (Friston et al. 1996). Any non-language related differences between language and control conditions, e.g. due to due to differential interactions between language areas and attentional, sensory, or motor networks can result in significant BOLD contrast. Considering the hemisphere-symmetric topography of the networks sub serving most brain functions(Salvador et al. 2005; Damoiseaux et al. 2006; Stark et al. 2008), a large portion of this confounding activity would be bilateral. This would cause a bias towards bilateral activity, pushing lateralization indices towards 0.

A second more speculative explanation is that while language is a-priori represented in both hemispheres, lateralization occurs as a result of either excitatory or inhibitory homotopic interhemispheric connections through the corpus callosum (Cook 1984; Gazzaniga 2000). While this interhemispheric interaction might result in one side being dominant, the non-dominant hemisphere might show a similar, yet attenuated activity pattern when the brain is processing language. This bilateral activity would also bias lateralization indices towards 0.

Regardless of which of these two explanations is valid, lateralization indices might benefit from a detailed homotopic mapping between hemispheres, where the focus is the activity difference between language activated voxels and their homologues, instead of the pooled number of activated voxels per hemisphere. Using the recently developed Cgrid-toolbox it is now possible to perform such a detailed homotopic mapping (Raemaekers et al. 2019). By using the Cgrid approach, we estimate a lateralization index based on the difference in activity between homotopic voxels in Broca’s and Wernicke’s areas. In addition we calculate the lateralization index using the more traditional approach. Results were compared to language lateralization as established with the WADA test, ECS or functional TransCranial Doppler sonography (fTCD).

## Methods

### Subjects

36 patients epilepsy patients were included in this study. The group consisted of ?? males (mean age ??, SD ??) and ?? females (mean age ??, SD). The patients were admitted to the University Medical Center Utrecht for epilepsy surgery in the period between 2006 and 2018. All patients gave written informed consent for participation.

### Scanning protocol

Patients were included over a 12 year period, during which substantial changes in scanner hardware and acquisition sequences occurred. Details on the acquisition parameters in each patient can be found in table 1 All functional images were acquired using a PRESTO protocol (Liu et al. 1993), and a T1-weigthted scan for anatomical reference was acquired in each patient.

### fMRI tasks

Data from two types of language paradigms were used, including the ‘Verb Generation’ (n=32) and ‘Picture Naming’ (n=29) tasks.

The verb generation task (VERBGEN) consisted of the presentation of written nouns, and subjects were instructed to silently think of a verb that can be associated to the noun. For example, the noun ‘ball’ can be associated with the verb ‘kick’. A baseline stimulus, consisting of squares/asterisks of similar size as the individual letters, and flashed with the same stimulus interval served as control condition.

During the picture naming task (PICNAM), images of common objects we presented, with subjects being instructed to silently name the object. The baseline stimulus included scrambled pictures of the same objects.

### WADA, ECS, and FTCD procedure

The WADA test consisted of an amobarbital injection in the internal carotid artery, which causes functional disruption of the ipsilateral cerebral hemisphere for 3–5 min. Meanwhile, the patient is asked to perform language tasks. If the patient can perform these without problems, language is localized on the contralateral to the injection. If the patient becomes aphasic, language is considered to be located to the injected hemisphere.

ECS was performed in patients in who a subdural electrode grid covering Broca’s and/or Wernicke’s area was temporarily implanted for epilepsy monitoring. During the testing protocol, pairs of adjacent electrodes were stimulated while the patient performed language tasks. If speech was disrupted by stimulation across any electrode pair, language was assumed to be located in the implanted hemisphere.

FTCD measures bilateral blood flow in the middle cerebral arteries using ultrasounds, which serves as a proxy of cerebral activity in Broca’s and Wernicke’s area (Woodhead et al. 2018). Changes in blood flow were measured during language tasks vs. baseline.

Details on which patients underwent which measurement can be found in table 1.

### MRI/fMRI Data processing

Functional MRI data were preprocessed using SPM12 (http://www.fil.ion.ucl.ac.uk/spm/). Preprocessing included realignment of the functional images and co-registration of the T1-weighted images to these images. The coregistered T1-weighted image was then submitted to the FreeSurfer recon-all pipeline, resulting in a surface reconstruction (Dale et al. 1999) and cortical labeling according to several atlases including amongst others the Desikan-Killiany-Tourville(DKT) (Klein and Tourville 2012) and Destrieux (Destrieux et al. 2010) atlas.

For homotopic mapping between the left and right hemisphere, we used the Cgrid-toolbox. The Cgrid-toolbox imposes a rectangular grid on predefined areas in the brain, and applying the same method to homologue areas results in 2D-matrix representations that are homotopically mapped onto each other. We created Cgrids of Broca’s area and its homologue, and of Wernicke’s area and its homologue.

The first step in generating Cgrids entails the extraction of patches of cortex based on the ROIs according to one of FreeSurfer cortical labeling schemes. These patches are subsequently flattened, and Cartesian grids are imposed by fitting polynomials to the borders of the patch (upper, lower, left, right), and subsequently interpolating these across the surface area or the patch. Cgrids can be composed of multiple concatenated subgrids, for each of which the borders have to be defined independently. The set of parameters for generating the Cgrids are shown in table 2A for Broca’s area, and 2B for Wernicke’s area, including also the x and y-resolution. Note that ROI definitions according to FreeSurfer’s Desikan-Killiany (Desikan et al. 2006) or DKT atlas did not allow for restricted inclusion of Wernicke’s area, so we used the more detailed Destrieux parcellation instead. An example of the imposed grids for Broca’s and Wernicke’s area for a single patient can be seen in figure 1.

**Figure 1:**
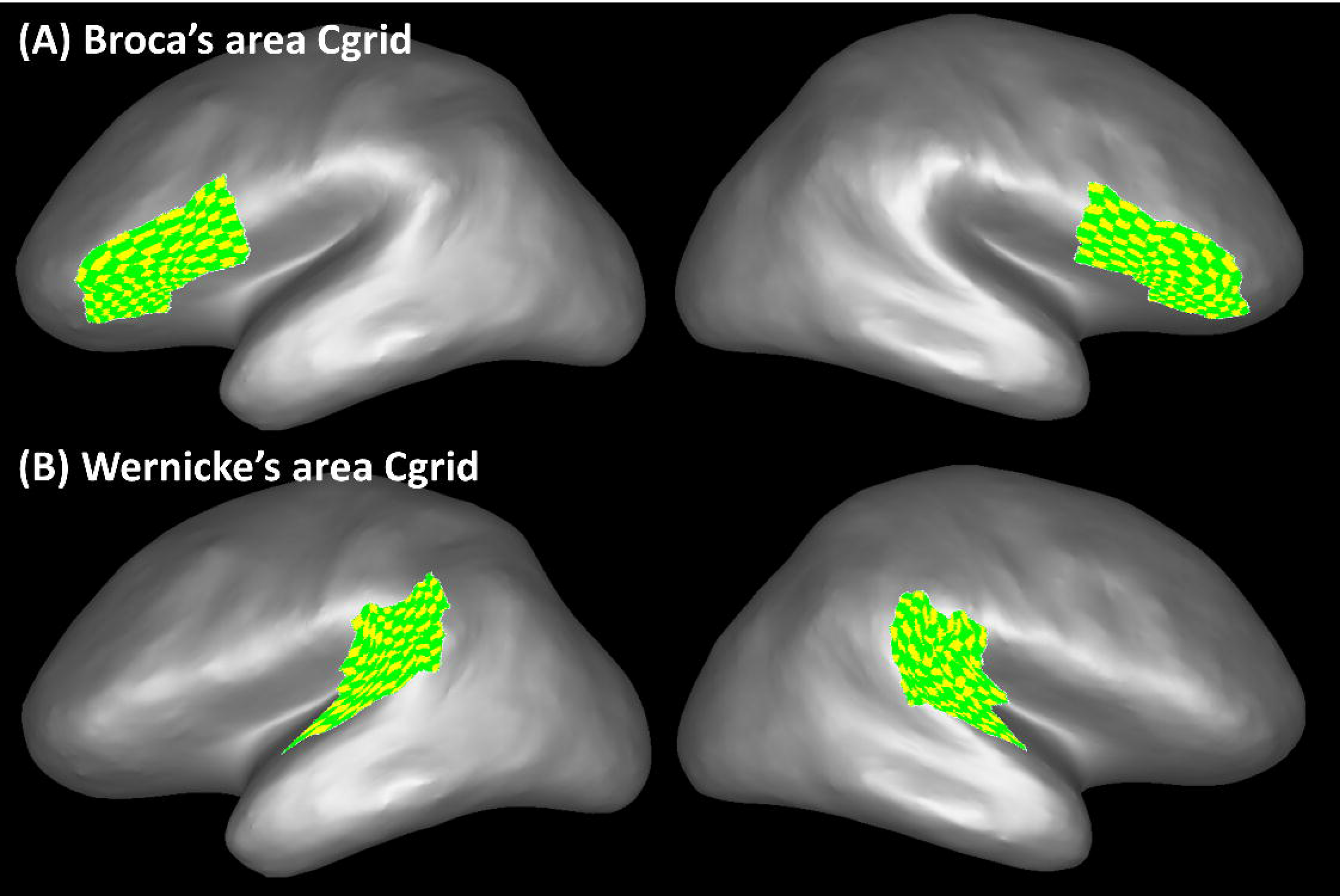
For a single patient, the grid structure projected on the cortical surface for (A) Broca’s area Cgrid and (B) Wernicke’s area Cgrid. Note that the resolution of the Cgrid created for generating the images was downsampled relative to the actually used Cgrids, to make the individual tiles better visible.

The preprocessed volumetric functional data of the two language tasks were mapped to the FreeSurfer surface reconstruction (specifically the middle grey matter layer) and smoothed across the surface using a 6 mm FWHM kernel. Subsequently, the functional surface data were mapped to the two language Cgrids and stored in 2 separate 4D Cgrid-nifti files for each subject. In the Cgrid-nifti format, the first 2 dimension represent the Cgrid x and y-coordinate, the third dimension represents the left or right hemisphere (1/2), and the fourth dimension the timeseries index. Results of right language lateralized patients according to the WADA test, ECS, of fTCD were flipped along the third dimension (left/right). In addition, for each subject and each of the 2 Cgrids a Cgrid-nifti file was created containing the sulcal depth to serve as background for functional results.

The Cgrid-nifti files containing the functional data were used as input in the first-level statistical analysis in SPM. The first-level design matrices included a single task-factor, explaining activity during the language task vs. the control conditions. A t-statistic was calculated representing more activity during the language than during the control conditions. The beta-maps resulting from the first-level analysis were included in second level one-sample t-tests for inspecting groupwise patterns of activity per hemisphere. A second paired-samples t-test was performed to inspect groupwise differences in the pattern of activity between the hemispheres, by contrasting the beta-maps and their versions flipped over the 3^rd^ dimension (left/right). Note that the Cgrid-nifti files are in a standard space, allowing direct comparisons across subjects and between hemispheres (Raemaekers et al. 2019).

The T-maps resulting from the first-level analysis were imported into MATLAB to calculate lateralization indices. For calculating traditional lateralization indices we used the formula:

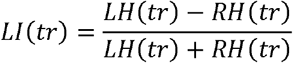

Where LH(tr) and RH(tr) are the number activated voxels in the left and right hemisphere respectively when using statistical threshold tr. For the novel approach the t-values in the right hemisphere were right hemisphere were subtracted from those in the left hemisphere. Then we applied the formula:

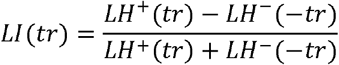

Where LH^+^ is the number of voxels more active in the left then right hemisphere, LH^−^ is the number of voxels more active in the right than left hemisphere, when using statistical threshold tr.

Note that lateralization indices strongly depend on the exact statistical threshold chosen for counting significantly activated voxels. Because of differences in the scanning protocol and intersubject variation in SNR of BOLD responses, we chose an approach where we include the 50% most activated voxels for calculating lateralization indices (Kristo et al. 2015). Differences in LI’s between the traditional and novel approach were calculated using a repeated measures GLM with 2 layers (traditional/novel approach) and four measures (VERBGEN-Broca, PICNAM-Broca, VERBGEN-Wernicke, PICNAM-Wernicke).

In addition, to avoid our results being specific to this single percentage threshold, we repeated the calculation across the full range of possible percentages (1%-100%) in steps of 1%. This provides eight 100-bin curves for each subject, i.e. for the novel and traditional approach, the VERBGEN and PICNAM, and the Broca’s and Wernicke’s Cgrids.

## Results

### WADA/ECS/fTCD results

All but one subject were left language lateralized according to the WADA test, ECS, or fTCD.

### Group-wise activity maps

Results of the main task effects are shown in Figure 2. Visual inspection of the results for the VERBGEN and PICNAM show the anticipated pattern, with more activity in the left than in the right hemisphere in both Broca’s and Wernicke’s area. The groupwise contrast between the hemispheres shows voxels more activated in the left than in the right hemisphere, but not vice versa (Figure 2) The number of significant voxels for the left/right contrast is however substantially smaller than the number activated for the main task effects in the left hemisphere.

**Figure 2:**
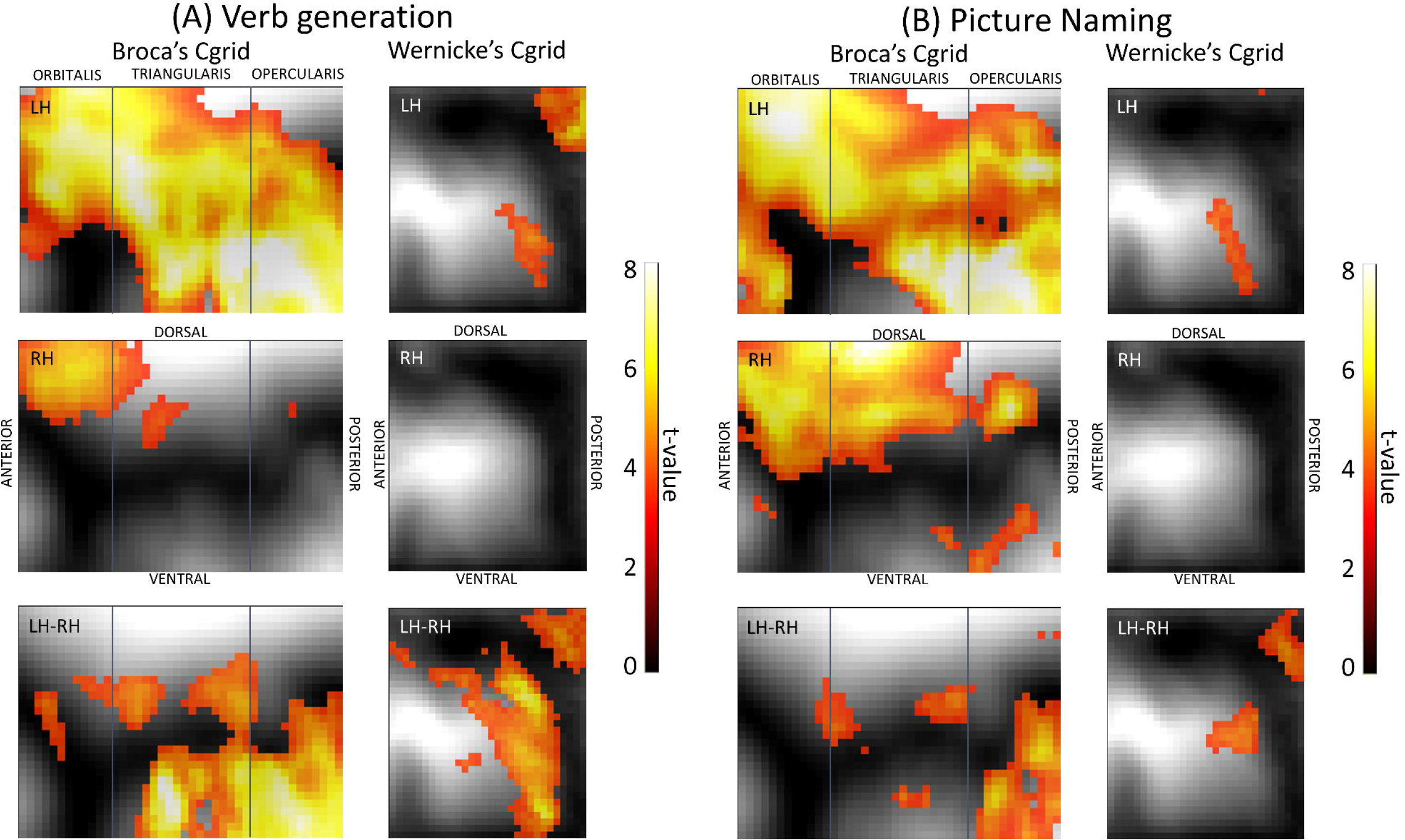
Groupwise activity during VERBGEN (A) and PICNAM (B) for Broca’s and Wernicke’s Cgrids. The top two panels show the activity in the left and right hemisphere respectively. The bottom panel shows the results for the difference in activity between the hemispheres (LH>RH). Significant voxels (p<0.001; uncorrected) are projected on the mean sulcal depth. There were no voxels where the t-value surpassed the negative statistical threshold. For the Broca’s area Cgrid, the borders between the individual subgrids are displayed.

### Lateralization indices

The multivariate test of the repeated measures GLM detected a significant effect of the approach for calculating LIs (F_(4,23)_=31.01;p<0.001). This effect was present for all combinations of task and area including VERBGEN in Broca’s area (F_(1,26)_=81.59;p<0.001), PICNAM in Broca’s area (F_(1,26)_=37.01;p<0.001), VERBGEN in Wernicke’s area (F_(1,26)_=26.79;p<0.001), and PICNAM in Wernicke’s area (F_(1,26)_=7.42;p=0.011). Visual inspection of the curves representing the mean LIs across the full range of included voxels showed a general pattern of increased LI’s using the novel approach, except for when including very few voxels (Figure 3). Effect sizes are further enhanced when observing the difference in LIs between the approaches (Figure 3).

**Figure 3:**
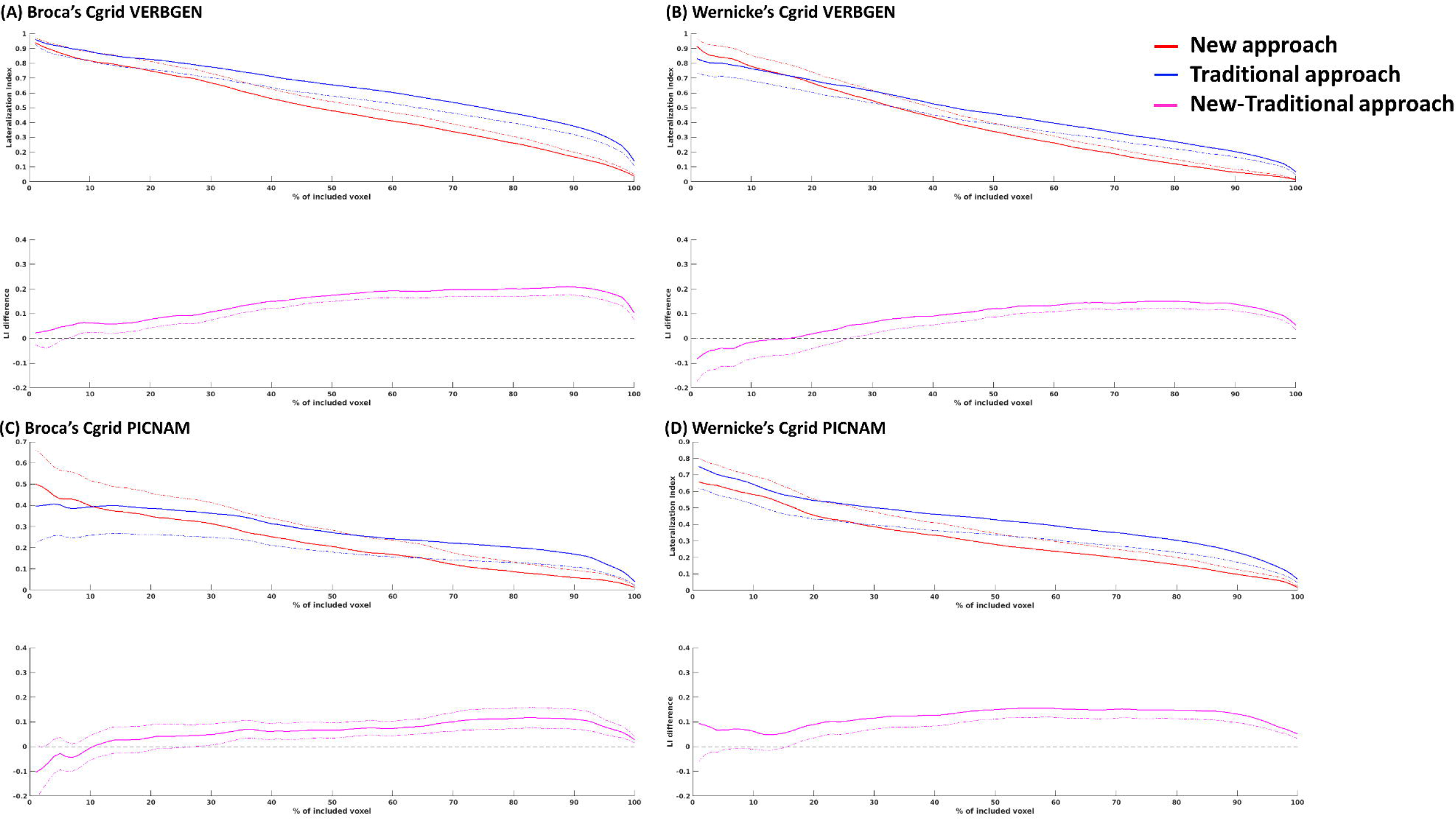
LIs with the inclusion of different percentages of most activated voxels for (A) Broca’s area Cgrid during VERBGEN, (B) Wernicke’s area Cgrid during VERBGEN, (C) Broca’s area Cgrid during PICNAM, and (D) Wernicke’s area Cgrid during PICNAM. The upper panel shows the LIs for the new and traditional approach. The lower panel shows the difference between the approaches. Dashed lines mark the 10% confidence interval (one-sided).

## Discussion

We calculated fMRI LIs using a traditional approach, meaning by comparison of the pooled amount of activated voxels per hemisphere, and contrasted this to a new approach based on the difference in activity between homotopic voxels. We found that using the new approach LIs became more outspoken in the direction of the language lateralization according to the WADA test, ECS, or fTCD.

We posed two possible mechanisms that might have caused the improvement. These were (1) the presence of confounding activity that is distributed in bilateral hemisphere symmetric activity as a result of imperfect task control (Friston et al. 1996), and (2) excitation or inhibition between of language areas and their homologues through homotopic connections through the corpus callosum (Cook 1984; Gazzaniga 2000). For the first case, bilateral symmetric activity that does not significantly differ between the left and right hemisphere would be expected. For the second case one would expect activity in the non- dominant hemisphere in areas that significantly differ between the hemispheres. For the first case we found evidence in the dorsal-anterior part of the Cgrid of Broca’s area (Figure 2). For the second case we found no evidence.

The control for confounding task activity of non-language related networks thus most likely plays a role in the found improvement when using the novel approach. In theory, this could also be achieved by an absolutely strict inclusion of language areas when calculating LIs. However the Cgrids that we used in this experiment already covered relatively restricted portions of known language related cortex. Using increasingly smaller areas would increase the risk of missing the language related activity altogether due to variations in functional topography across subjects (Xiong et al. 2000).

One important disadvantage if the novel approach is that it requires substantial additional processing relative to the traditional approach, including the generation of surface reconstructions and subsequently the Cgrid representations. While most of processing steps can be performed fully automated, making application easily accessible, the generation of surface reconstructions more or less assumes intact cortex. This implies possible complications in subjects with large neurological abnormalities. While this is most likely not an issue for most epilepsy patients undergoing surgery, as their T1-weighted images are often (close to) similar to that of healthy controls. The situation can be very different when considering tumor patients, where the generation of surface reconstructions and Cgrids can be seriously complicated.

The fMRI data used in the current study was obtained over a long time period during which various changes in scanner hardware and scanning protocols occurred. We accepted these variations to maximize patient inclusion. The variations in the acquisition in all likelihood produced an increase in the between-subject variation of fMRI signal to noise ratios, and subsequently an increase in the variation in lateralization indices across subjects. While not optimal, the impact on the paired comparison between the old and the new approach might be relatively limited, as both approaches used the same data. We thus believe that the strategy to maximize patient inclusion resulted in the optimal statistical power for this study.

All but one patient in this sample were left language lateralized according to the WADA test, ECS, or fTCD. Although the only right lateralized patient did show an improvement, and there is no a-priori reason to believe that matters will be different for left language lateralized subjects, this results can in principle not be generalized. The obtained results thus need to be confirmed in a sample with only right language lateralized subjects.

For homotopic mapping between the hemispheres we used the Cgrid approach. Other methods have been proposed for achieving homotopic mapping between hemispheres, including e.g. by registration to a symmetric MNI template (Zuo et al. 2010). Although we did not directly assess the performance of this alternative, we believe that our method has the advantage of exploiting the inherent benefits of surface based registration (Ghosh et al. 2010), resulting in a closer correspondence between homologue voxels.

The current approach might also prove to be relevant for reducing the number of false positive sites when comparing fMRI language activity to intra-operative ECS. When assuming ECS as gold standard, fMRI produces relatively few type II errors, but a large proportion of type I errors (Giussani et al. 2010). The results of this study suggest that this situation might be improved by comparing ECS to the difference in activity between pairs of homotopic voxels. Note however that while this might increase specificity, the caveat is that some amount of spatial smoothing is necessary to compensate for errors in the mapping between the hemispheres. Depending on the amount of smoothing necessary, the spatial resolution of the fMRI results will be affected. In this study we used 6 mm FWHM Gaussian smoothing across the surface,

In summary, we propose an alternative way of calculating lateralization indices that controls for confounding brain activity. This control brings LIs according to fMRI in closer alignment with language lateralization as determined by the WADA test, ECS, or fTCD. These results improve the prospects of fMRI replacing the more invasive WADA test in the future.

## Legends

Table 1: fMRI scanning parameters and WADA/ECS/fTCD results for each patient.

Table 2: Parameters used for creating the Cgrids of (A) Broca’s area and (B) Wernicke’s area.

