## Supplementary material for "Enhancing fMRI language lateralization indices by homotopic mapping": table 1

| **Patients** | **Year** | **MR Field strength**  **(T)** | **VERBGEN**  **Scantime**  **Dynamic (s)** | **PICNAME**  **Scantime**  **Dynamic (s)** | **VERBGEN**  **# of acquired volumes** | **VERBGEN**  **# of acquired volumes** | **WADA** | **FTCD** | **ECS** |
| --- | --- | --- | --- | --- | --- | --- | --- | --- | --- |
| Aart | 2006 | 1.5 | 2.416 | 1.489 | 120 | 320 | LH |  |  |
| Ahma | 2014 | 3 | 0.608 | 0.608 | 531 | 401 |  |  | LH |
| Arno | 2009 | 3 | 0.608 | 0.608 | 486 | 401 |  |  | LH |
| Beil | 2018 | 3 | 0.608 | 0.608 | 486 | 401 |  | LH |  |
| Bred | 2015 | 3 | 0.608 | 0.608 | 486 | 401 |  |  | LH |
| Brem | 2009 | 3 | 0.608 | 0.608 | 482 | 401 | LH |  |  |
| Bunn | 2016 | 3 | 0.608 | 0.608 | 486 | 401 |  | LH |  |
| Esse | 2012 | 3 | 0.608 | 0.608 | 486 | 401 | LH |  |  |
| Giel | 2014 | 3 | 0.608 | 0.608 | 482 | 401 |  |  | LH |
| Gord | 2008 | 3 | 0.608 | 0.608 | 486 | 401 | LH |  |  |
| Graf | 2011 | 3 | 0.608 | 0.608 | 482 | 401 | LH |  |  |
| Groe | 2010 | 3 | 0.608 | 0.608 | 482 | 401 | LH |  |  |
| Groo | 2008 | 3 | 0.608 | 0.608 | 486 | 801 | LH |  |  |
| Guij | 2011 | 3 | 0.608 | 0.608 | 486 | 401 |  |  | LH* |
| Habe | 2010 | 3 | 0.608 | 0.608 | 482 | 401 | LH |  |  |
| Heem | 2006 | 3 | 2.416 | 1.489 | 120 | 320 | LH |  |  |
| Hube | 2013 | 3 | 0.608 | 0.608 | 482 | 401 |  |  | LH |
| Jans | 2008 | 3 | 0.608 | - | 486 | - | RH |  |  |
| Maan | 2007 | 1.5 | 2.416 | 1.489 | 120 | 320 | LH |  |  |
| Mang | 2007 | 1.5 | 2.416 | 1.489 | 120 | 320 | LH |  |  |
| Mels | 2010 | 3 | 0.608 | 0.608 | 482 | - | LH* |  |  |
| Pol_ | 2007 | 1.5 | 2.416 | 1.489 | 120 | 320 | LH |  |  |
| Rech | 2013 | 3 | 0.608 | 0.608 | 482 | 401 |  |  | LH* |
| Rooi | 2010 | 3 | 0.608 | 0.608 | 482 | 401 | LH* |  |  |
| Ruit | 2009 | 3 | - | 0.608 | - | 401 |  |  | LH |
| Siez | 2010 | 3 | 0.608 | 0.608 | 486 | 401 | LH |  |  |
| Sijm | 2008 | 3 | 0.608 | 0.608 | 486 | 401 | LH |  |  |
| Smit | 2006 | 1.5 | 2.416 | 1.489 | 120 | 320 | LH |  |  |
| Spoo | 2008 | 3 | 0.608 | 0.608 | 486 | 401 |  |  | LH* |
| Ve3_ | 2013 | 3 | 0.608 | 0.608 | 482 | 401 | LH |  |  |
| Viss | 2010 | 3 | 0.608 | - | 482 | - | LH |  |  |
| Vled | 2016 | 3 | 0.608 | - | 486 | - |  |  | LH |
| Weve | 2007 | 1.5 | 2.416 | 1.489 | 120 | 320 |  |  | LH* |
