## Supplementary material for "Enhancing fMRI language lateralization indices by homotopic mapping": table 2

| **(A) BROCA** | Subgrid1 | Subgrid2 | Subgrid3 |
| --- | --- | --- | --- |
| Included area | Parsorbitalis | Parstriangularis | Parsopercularis |
| Upper border | Parstriangularis | Parsopercularis | Precentral |
| Lower border | Lateral  orbitofrontal | Parsorbitalis | Parstriangularis |
| Right border | Insula | Insula | Insula |
| Left border | Rostral  middlefrontal | Rostral  middlefrontal | Rostral  middlefrontal  Caudal  middlefrontal |
| x-resolution | 30 | 30 | 30 |
| y-resolution | 12 | 18 | 12 |

| **(B) WERNICKE** | Grid |
| --- | --- |
| Included area | A. G Pariet Inf - Supramar B. G Temp Sup - G T Transv C. G Temp Sup - Plan Tempo D. Lat Fis - Post E. S Temporal Transverse |
| Upper border | **A** - S Postcentral  **A** - G Pariet Sup  **A** - G Postcentral |
| Lower border | **A** - G Temp Sup Lateral  **B** - G Temp Sup Lateral  **C** - G Temp Sup Lateral  **E** - G Temp Sup Lateral |
| Right border | **A** - G Pariet Inf Angular  **A** - S Intrapariet & P Trans  **A** - S Temporal Sup |
| Left border | **A** - G & S Subcentral  **B** - S Circular Insula Sup  **B** - S Circular Insula Inf  **B** - G Temp Sup Plan Polar  **D** - G Ins LG & S Cent Ins |
| x-resolution | 30 |
| y-resolution | 35 |
